# Electrophysiological characterization of AtAMT1;4, an extraordinarily high affinity ammonium transporter from *Arabidopsis thaliana*

**DOI:** 10.1101/820712

**Authors:** Nino Bindel, Benjamin Neuhäuser

**Affiliations:** Institute of Crop Science, Nutritional Crop Physiology, University of Hohenheim, Fruwirthstr. 20, D-70593 Stuttgart, Germany

**Keywords:** ammonium transport, plant nitrogen nutrition, high affinity, electrophysiology

## Abstract

In plants high affinity transport proteins mediate the essential transport of ammonium across membranes. In *Arabidopsis thaliana* six of these <u>AM</u>monium <u>T</u>ransporters (AMTs) are encoded on the genome. All of these show a unique expression pattern. While most AMTs are highly expressed in the root, AtAMT1;4 expression is limited to the pollen grains and the pollen tube. Here, we addressed the transport characteristics of AtAMT1;4 in the heterologous *Xenopus laevis* oocytes system. Two electrode voltage clamp measurements tagged AtAMT1;4 as an electrogenic high affinity ammonium transporter. The transport was saturable and showed extraordinarily high affinity for ammonium with a K_m_ value lower than 10 µM.

---

To meet the requirements for fast and sufficient nitrogen uptake and distribution within the organism, several distinct transport proteins for ammonium have evolved in the model plant *Arabidopsis thaliana* ^1^. Six of these <u>AM</u>monium <u>T</u>ransporters (AMTs) are encoded on the Arabidopsis genome, all of them showing diverse localization pointing to specialized functions in the process of ammonium distribution throughout the plant ^2^. While AtAMT1;5 expression and activity is negligible, three of the high affinity AMT1 transporters are mainly located in the roots to mediate the transport of ammonium from the soil to the vasculature ^1,2^. AtAMT1;1 and AtAMT1;3 are located in the plasma membrane of the epidermal and cortical cell layer of the root ^3,4^. AtAMT1;2 localization is restricted to the cortical and endodermal root cells ^1^. The localization and high affinity of AtAMT1;1 (In oocytes: K_m_ = 2.7 µM ^3^; in plants: K_m_ = 50.0 µM ^2^) and AtAMT1;3 (In oocytes: K_m_ = 129 µM ^5^; in plants: K_m_ = 60.5 µM ^2^) suggest a function of these two transporters in the direct primary ammonium uptake from the soil. Further transport of ammonium into the vascular tissue depends on transfer across the casparian strip. AtAMT1;2 with lower affinity but localization in the endodermis was proposed to mediate the uptake into the endodermal cell layer, where apoplastic flow is blocked by the casparian strip ^1^. Indeed it was shown that nitrogen allocation to the shoot is AtAMT1;2 dependent ^6^.

AtAMT2, the sole member of the AMT2 subfamily in Arabidopsis, is strongly expressed in the root as well. While expression in the young root parts can be seen in the trichoblasts and root-hair ^7^, localization in the older root parts shifts to a preferential expression in the pericycle ^8^. Expression in the trichoblasts implies a function in primary ammonium uptake while expression in the pericycle was connected with a function in long distance ammonium transfer to the shoot ^7,8^. In strong contrast to these transporters, AtAMT1;4 expression is exclusively found in the pollen grains and the pollen tube ^9^. This localization excludes a function in primary ammonium uptake. Next to AtAMT1;4 several transporters for organic nitrogen forms are highly expressed in the pollen, suggesting their role in nitrogen loading into the pollen ^10,11^.

General ammonium transport function of AtAMT1;4 was previously shown by complementation of an ammonium transporter deficient yeast. Further tissue independent overexpression of AtAMT1;4 conferred high affinity ammonium uptake capacity to uptake deficient *Arabidopsis* plants ^9^.

In this work we readdressed the ammonium transport function of AtAMT1;4 in the background free *Xenopus leavis* oocyte system. We compared AtAMT1;4 with AtAMT1;1 mediated transport. Ammonia transport by AtAMT1;4 was proton coupled and elicited strong inward currents. The net ammonium transport by AtAMT1;4 was saturable with a strongly potential dependent affinity.

## Results

### Proton coupled ammonium transport by AtAMT1;1 and AtAMT1;4

Oocytes expressing AtAMT1;1 and AtAMT1;4 were subjected to ammonium containing recording solution. Ammonium in a concentration of 300 µM induced strong inward currents with a reversal potential between +60 mV and +80 mV (Abb. 1). In non-injected or water-injected controls ammonium did not elicit ionic inward currents. The currents induced by both transporters were voltage dependent and declined with decreasing membrane potential.

### AtAMT1;4 shows extraordinarily high affinity for ammonium

When expressed in oocytes both transporters elicited ammonium concentration dependent currents (Fig. 2 A/C). Those currents saturated in a membrane potential dependent manner. While half maximal currents of AtAMT1;1 at −120 mV were reached at K_m-120mV_ = 16 µM, AtAMT1;4 mediated currents saturated with K_m-120mV_ = 6.5 µM. The affinity for ammonium was strongly potential dependent. K_m_ values increased at less negative membrane potential. The K_m_/potential relationship is shown in Fig. 2 B/D. This relationship was fitted as an exponential curve with the following equation: K_m_(δ) = K_m (0 mV)_ X exp (δ X *e* X V / *k* X T). δ gives the fractional electrical distance which indicates the point in the membrane electric field where the ion is first bound to the protein. This δ value was almost similar for both transporters with δ_AtAMT1;1_ = 0.357 and δ_AtAMT1;4_ = 0.372. Therefore, the binding site for ammonium lies deep inside the transporter pores after crossing approximately 36 % or 37 % of the membrane electric field.

## Discussion

Like the other AtAMT1 transporters, which are involved in high affinity primary ammonium uptake into the root, the pollen localized AtAMT1;4 mediates electrogenic ammonium transport when subjected to 300 µM ammonium chloride. Therefore, either the ion is directly transported or ammonium is recruited at a binding-side approximately 37 % inside the membrane electric field (shown by the δ values) and then deprotonated during the further transport process. Binding of ammonium followed by deprotonation was proposed based on molecular dynamic transport simulation ^12,13^ and has recently been suggested to be a general feature of proteins from the AMT (<u>AM</u>monium <u>T</u>ransporter)/ Mep (<u>Me</u>thylammonium <u>P</u>ermease)/ Rh (<u>Rh</u>esus) family ^14^. A binding side located deeply inside the transporter pore is in accordance with calculations for other plant AMT1 proteins ^1,15^. In AmtB from *E. coli* a phenylalanine gate blocking the open pore and preventing further uncontrolled ion flux is located approximately 40 % inside the membrane electric field ^16–18^. These Phenylalanine residues are highly conserved in AMT/MEP/Rh proteins and are directly involved in blocking ammonium ion flux through the central pore (Ganz et al. under review).

The ammonium transport of both AMT1s tested, was stable and saturable. Both transporters showed high affinity in the micro molar range. For AtAMT1;1 this high affinity is easily explained since plants are always competing for sufficient ammonium uptake from the soil with other plants or microorganisms. The high ammonium affinity of AtAMT1;1 was shown before ^3^ but here we were able to show much higher ammonium induced currents (Fig. 1 and 2). In the previous study AtAMT1;1 currents reached about 30 nA when oocytes were supplied with 300 µM ammonium. The ammonium currents reported here were much more stable and more than ten times higher (Fig.1).

**Fig 1:**
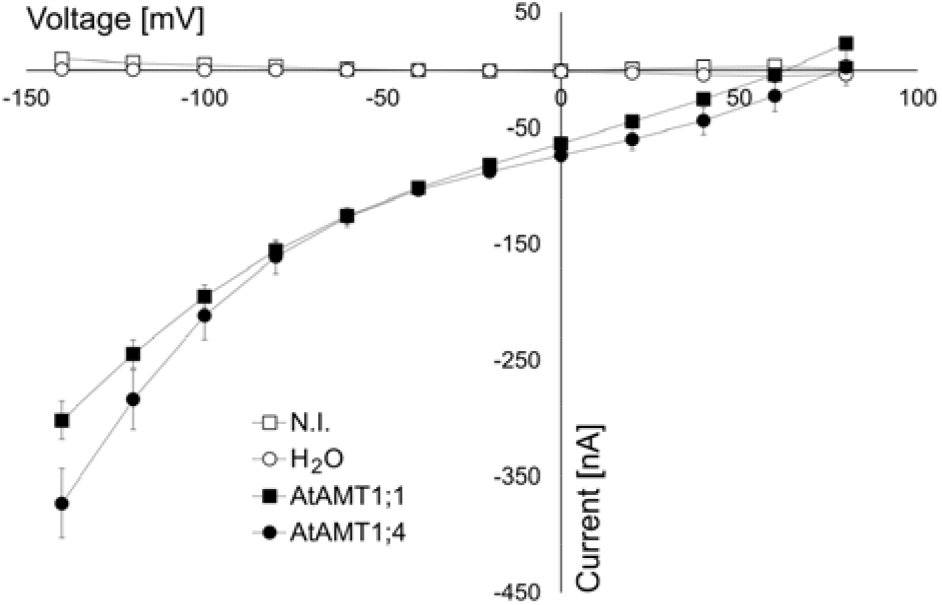
Mean current / voltage relationship of the AtAMT1;1 and AtAMT1;4 mediated currents elicited by 300 µM ammonium. Oocytes expressing AtAMT1;1 or AtAMT1;4 were subjected to recording solution containing 300 µM ammonium. Both proteins mediated stable inward directed currents that decreased with decreasing membrane potential. Data is given as means (± SE), n = 4.

**Fig 2:**
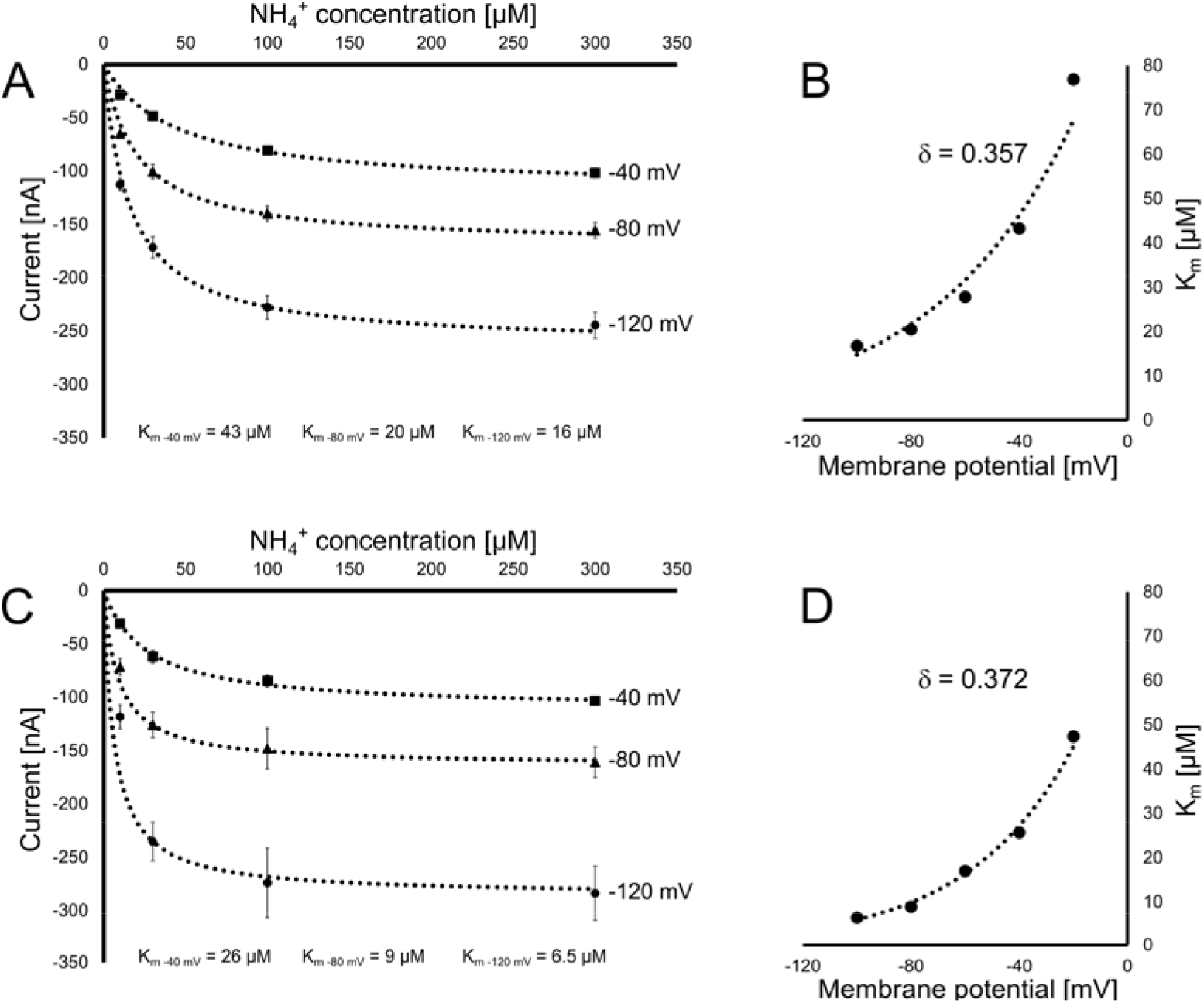
Transport kinetics of AtAMT1;1 and AtAMT1;4. When expressed in oocytes both transporters elicited ammonium dependent currents [**(A)** AtAMT1;1 and **(C)** AtAMT1;4]. The currents saturated at increasing concentrations. This transport kinetics was strongly membrane potential dependent and affinity increased (K_m_ decreased) with decreasing membrane potential [**(B)** AtAMT1;1 and **(D)** AtAMT1;4]. Data is given as means (± SE), n ≥ 4.

We here report the first quantitative transport analysis of AtAMT1;4 in a background free heterologous system. AtAMT1;4 mediated currents were stable and saturated in a membrane potential dependent manner, as well. AtAMT1;4 exhibited an extremely high ammonium affinity which was superior to AtAMT1;1 affinity (Fig. 2). Similarly high K_m_ values for AtAMT1;4 had been shown by complementation analysis of an ammonium transporter deficient *Arabidopsis* line ^9^. The physiologic function of this high affinity can only be speculated on. The expression in the young microspores suggests that ammonium transport is needed to supply the microspore with nitrogen for its further development ^9^. While this may not explain the extraordinarily high substrate affinity it allows us to speculate that free ammonium concentrations in the surrounding anther tissue are minor.

This study confirmed high affinity net ammonium transport by AtAMT1;1 and AtAMT1;4. The reported currents for AtAMT1;1 were more stable than in previous studies. For AtAMT1;4 we show the first characterization of transport in a background free heterologous system.

## Experimental procedures

### Construct preparation

The coding sequence of AtAMT1;4 (At4g28700) was amplified from genomic Col-0 DNA. Primers sequences for amplification included restriction enzyme cut sites for XbaI and NcoI. (Primer sequences: AtAMT1;4-XbaI-Fw: ATAtctagaATG GCGTCGGCTCTCTCTT AtAMT1;4-NcoI-Rv: ATTccatggTCAAACACCTACATTGGGATCATTATC) The PCR product as well as the pOO2 vector were both XbaI/NcoI digested, cleaned by gel-electrophorese and T4 ligated. Correct insertion of the AtAMT1;4 CDS into pOO2 was tested by Sanger-sequencing. AtAMT1;1-pOO2 was present in the lab ^3^.

### Preparation of cRNA and injection of oocytes

Dissected and preselected *Xenopus laevis* oocytes were obtained from Ecocyte Bioscience (Dortmund, Germany). Oocytes were stored in ND96 solution (96 mM NaCl, 2 mM KCl, 1 mM MgCl_2_, 1.8 mM CaCl_2_, 2.5 mM sodium pyruvate, 5 mM HEPES adjusted to pH of 7.4 by NaOH) at 4°C prior to injection. cRNA was produced from MluI linearized and phenol-chloroform purified pOO2 plasmids containing the CDS of AtAMT1;1 or AtAMT1;4 using the mMESSAGE mMACHINE™ SP6 Transcription Kit (Life Technologies GmbH, Darmstadt; Germany) following the manufactures instructions. 50 nl cRNA with a concentration of 400 ng/µl were injected into each oocyte. Oocytes were kept in ND96 for 3 days at 18 °C before electrophysiological measurements were performed.

### Electrophysiological measurements

Electrophysiology was performed in a small recording chamber (as described before ^3^) containing Choline-Cl recording solution (110 mM choline chloride, 2 mM CaCl_2_, 2 mM MgCl_2_, 5 mM MES, pH adjusted to 5.5 with Tris). Variable ammonium concentrations were added as NH_4_Cl salt. Currents without added ammonium were subtracted at each voltage. The concentration dependence of currents was fitted using the following equation: I = I_max_ / (1 + K_m_ / c), with I_max_ (maximal current at saturating concentration), K_m_ (substrate concentration permitting half-maximal currents), c (experimentally used ammonium concentration). The voltage dependence δ of the K_m_ was calculated using the following equation: K_m_(δ) = K_m (0 mV)_ X exp (δ X *e* X V / *k* X T), with δ (fractional electrical distance), *e* (elementary charge), V (membrane potential), *k* (Boltzmann’s constant), and T (absolute temperature).

## Acknowledgments

We thank Uwe Ludewig for discussion.

## Author contributions

**N.B.** and **B.N.** performed the experiments. **B.N.** analysed the data. **B.N.** planed the experiments and wrote the manuscript. **B.N.** is the corresponding author.

